# Subinhibitory antibiotic concentrations promote the excision of a genomic island carried by the globally spread carbapenem-resistant *Klebsiella pneumoniae* ST258

**DOI:** 10.1101/2023.08.10.552780

**Authors:** Alejandro Piña-Iturbe, Guillermo Hoppe-Elsholz, Isidora D. Suazo, Alexis M. Kalergis, Susan M. Bueno

## Abstract

The ICEKp258.2 genomic island (GI) has been proposed as an important factor for the emergence and success of the globally spread carbapenem-resistant *Klebsiella pneumoniae* sequence type (ST) 258. However, a characterization of this horizontally acquired element is lacking. Using bioinformatic and experimental approaches, we found that ICEKp258.2 is not confined to ST258 and ST512 but also carried by ST3795 strains and emergent invasive multidrug-resistant pathogens from ST1519. We also identified several ICEKp258.2-like GIs spread among different *K. pneumoniae* STs, other *Klebsiella* species, and even other pathogen genera, uncovering horizontal gene transfer events between different STs and bacterial genera. Also, in agreement with the origin of ST258 from ST11, the comparative and phylogenetic analyses of the ICEKp258.2-like GIs suggested that ICEKp258.2 was acquired from an ST11 strain. Importantly, we found that subinhibitory concentrations of antibiotics used in treating *K. pneumoniae* infections can induce the excision of this GI and modulate its gene expression. Our findings provide the basis for the study of ICEKp258.2 and its role in the success of *K. pneumoniae* ST258. They also highlight the potential role of antibiotics in the spread of ICEKp258.2-like GIs among bacterial pathogens.

## INTRODUCTION

Carbapenem-resistant *Klebsiella pneumoniae* (CR-*Kp*) is one of the major contributors to the global deaths caused by antibiotic-resistant bacteria and, therefore, is a current threat to public health which can cause hospital-acquired pneumonia, urinary tract infections, and bacteriemia in susceptible individuals, leading to elevated treatment costs, increased hospitalization times and higher mortality rates (1–3). The clonal complex 258, which includes the sequence-types (ST) 11 and 258 among other closely related STs, has received particular attention due to their worldwide distribution, high prevalence, and its role in the global dissemination of carbapenemases such as those encoded by the highly prevalent *bla*_KPC-2_ and *bla*_KPC-3_ genes (4–6). Over the past two decades, the ST258 has become one of the most successful clones of CR-*Kp* found in healthcare settings worldwide, especially in North America and Europe (7–9). However, the factors explaining its successful spread are still unknown, and their identification is considered an important question with impact in public health (10, 11).

Comparative genomics and phylogenetic analyses have shown that the epidemic CR-*Kp* ST258 emerged as the result of several modifications at the genetic and genomic scale, some of which may be involved in the success of this pathogen. Analysis of the single-nucleotide polymorphisms distribution between ST11, ST442, and ST258 strains revealed that the ancestral ST258 emerged from a large recombination event that originated its hybrid chromosome comprised of ≈80% from ST11 and ≈20% (1.06 Mbp) from ST442 (12). This 1.06 Mbp recombination region includes the *tonB* allele, which differentiates ST258 (*tonB79*) from ST11 (*tonB4*), and the *cps* locus (capsule type *wzi154*/KL107), involved in the synthesis of the capsular polysaccharide (12). Phylogenetic analyses showed that after the main chromosomal recombination event, the ancestral ST258 acquired mutations in the *marR* and *ompK35* genes, which resulted in a Phe34>Ser34 substitution in the transcriptional regulator MarR and a truncated porin OmpK35; the MarR mutation is hypothesized to impact the global metabolism in ST258, and truncation of OmpK35 has been shown to increase the resistance to different antibiotics, including carbapenems (7, 13, 14). In addition to these mutations, the ancestral ST258 acquired the ICEKp258.2 genomic island (GI), integrated into the 1.06 Mb recombination region, followed by the replacement of the region carrying the *cps* locus (≈52 kbp) with the corresponding region of an ST42 strain (capsule type *wzi29*/KL106), resulting in the emergence of the Clade I ST258 (clade II ST258 harbors the original *wzi154*/KL107 capsule type) (12, 13, 15). Finally, events of horizontal acquisition and vertical transmission in both clade I and clade II ST258 strains shaped the *bla_KPC_*-carrying plasmid pool that characterizes the epidemic CR-*Kp* ST258 strains, which is diverse and include transferable plasmids from different incompatibility groups (7, 10, 13). Since antibiotic resistance, including the carry of the *bla_KPC_* genes, is not restricted to ST258, and strains of this ST show a relatively low virulence in animal models or harbor virulence factors inconsistently, the role of other factors besides multidrug resistance and variable virulence determinants are being the subject of hypotheses and research aiming to understand the success of CR-*Kp* ST258 (10, 11, 16, 17).

It has been hypothesized that the acquisition of the ICEKp258.2 GI, which took place after the emergence of ST258 but before its worldwide dissemination, has played a role in the spread of this pathogen (12). This hypothesis is based mainly on the findings that ICEKp258.2 seems to be restricted to ST258 and carries the genes predicted to encode a putative type-III restriction-modification (RM) system and putative type-IV-pili (T4P)-related proteins (12, 13, 18). It has been postulated that the T4P-related proteins may contribute to bacterial adherence to surfaces or the uptake of foreign DNA. At the same time, the Type-III RM system may restrict the acquisition of foreign genetic elements, thus shaping the plasmid pool associated with ST258 (12). However, despite its possible role in the success of CR-*Kp* ST258, characterization of this GI is lacking. In previous work from our laboratory, we found that ICEKp258.2 can be excised from the bacterial chromosome and is a member of a widespread family of excisable GIs found in different species from the order Enterobacterales, the Enterobacterales-associated ROD21-like (EARL) family (19).

Here, we expand our analysis, providing a comprehensive characterization of the gene content, conservation, host distribution, and phylogenetic relationships of ICEKp258.2, as well as the effect of subinhibitory concentrations of clinically relevant antibiotics, including carbapenems, on its integration/excision state and gene expression. We found evidence indicating that ICEKp258.2 was likely acquired from an ST11 strain, and that active horizontal transfer and modular recombination events are disseminating closely related GIs among different species of pathogenic bacteria. Moreover, we found a positive correlation between antibiotic potency and the excision frequency of ICEKp258.2, unveiling a potential role of antibiotics in the spread of the ICEKp258.2-like GIs among bacterial pathogens.

## RESULTS

### The ICEKp258.2 genomic island shares a conserved backbone with the EARL GIs and forms a circular element upon excision

Originally identified through comparative genome analyses between *K. pneumoniae* strains from different STs and ST258, the ICEKp258.2 GI was found as a low-GC (37.1%) horizontally acquired element, integrated into the 1.06 Mbp recombination region in an Asn-tRNA-encoding gene and carried exclusively by ST258 strains and its single locus variant (SLV) ST512 (12, 15, 18) (**Fig. 1A, B**). However, a more comprehensive characterization was lacking. Therefore, we started by examining the ICEKp258.2 genetic context and gene content in the prototypical strain NJST258_2.

**Figure 1.**
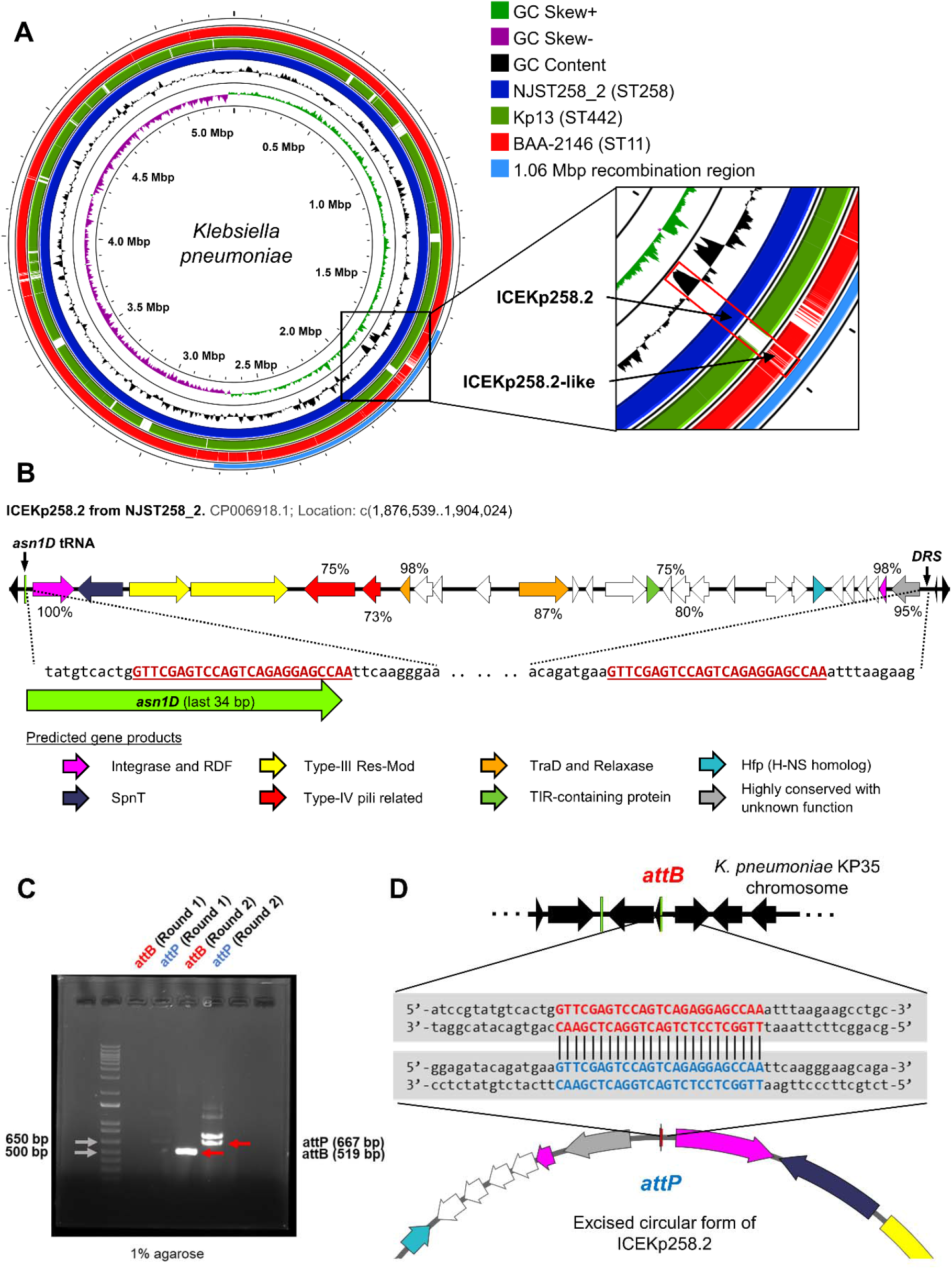
The ICEKp258.2 share a conserved backbone with the EARL GIs and forms a circular element upon excision. A) Comparison of representative CR-*Kp* genomes from strains BAA-2146 (ST11; CP006659.2), Kp13 (ST442; CP003999.1), and NJST258_2 (ST258; NZ_CP006918.1). The zoomed-in panel shows the location of the ICEKp258.2 GI within the 1.06 Mbp recombination region of NSJT258_2, its absence in Kp13 and an ICEKp258.2-like GI in BAA-2146. **B)** Genetic organization and integration site of the ICEKp258.2 GI from strain NJST258_2. The colors represent the predicted gene products and the numbers indicate the presence of homologous genes among the EARL GIs identified in (19). The direct-repeated sequences (DRS) located at both ends of the GI are also indicated. **C)** Agarose gel showing the amplification products of primers targeting the insertion (*attB*) site and the circularized form (*attP*) of the ICEKp258.2 GI from *K. pneumoniae* strain KP35 (ST258). Visible bands were obtained after a second round of PCR using the same primers and 2 µL of the reaction mix from the first round as template. **D)** Nucleotide sequences determined by Sanger sequencing of the purified PCR products in **C**. The *attB* and *attP* core sequences are highlighted in red and blue font.

ICEKp258.2 was found integrated at the 3’-end of *asn1D* and flanked by two perfect direct repeated sequences of 25 bp (GTT CGA GTC CAG TCA GAG GAG CCA A), which delimited the 27,486 bp GI (**Fig. 1B**). *asn1D* is one of the four identical Asn-tRNA-encoding genes found in the *Klebsiella* genome, which represent an integration hotspot for different island families, including the EARL GIs (19–21). Most of the 28 open reading frames (ORFs) carried by ICEKp258.2 (19/28) are predicted to encode hypothetical proteins of unknown function (**Table 1**). However, all ORFs were shared by at least another EARL GI (**Supplemental Table S1**). The most conserved ORFs, present in ≥70% of the 55 previously identified EARL GIs, correspond to those predicted to encode the proteins involved in the excision/integration-and conjugation-related functions of the GI (i.e., an integrase and recombination directionality factor, and TraD and putative relaxase from MOB_Q_ family), as well as the pilus-related proteins and three hypothetical unknown proteins (**Fig. 1B**). The ORFs encoding the restriction and methylation subunits of a putative type-III RM system had low conservation among the EARL GIs (36 and 33%, respectively). However, this results from both genes being highly prevalent only in one EARL family clade, which includes ICEKp258.2 and GIs harbored by a high diversity of bacterial species (**Supplemental Table S1**) (19).

**Table 1.** Annotation, location, and predicted products of the ORFs within ICEKp258.2 from *Klebsiella pneumoniae* ST258 strain NSJT258_2

| ORF | Locus tag | GenBank RefSeq<br>annotation | Location | ORF<br>length<br>(bp) | Protein<br>length<br>(aa) |
| --- | --- | --- | --- | --- | --- |
|  | KPNJ2_RS09445 | tRNA-Asn | c1904001..1904076 | 76 | -- |
| 1 | KPNJ2_RS09440 | integrase arm-type<br>DNA-binding domain-containing protein | c1902546..1903820 | 1275 | 424 |
| 2 | KPNJ2_RS09435 | hypothetical protein<br>(SpnT)<sup>a</sup> | 1901086..1902474 | 1389 | 462 |
| 3 | KPNJ2_RS09430 | site-specific DNA-methyltransferase | c1,899,042..1,900,892 | 1851 | 616 |
| 4 | KPNJ2_RS09425 | DEAD/DEAH box<br>helicase family<br>protein | c1,896,072..1,899,032 | 2961 | 986 |
| 5 | KPNJ2_RS27875 | type II secretion<br>system protein | 1,894,050..1,895,609 | 1560 | 519 |
| 6 | KPNJ2_RS09415 | type 4 pilus major<br>pilin | 1,893,277..1,893,846 | 570 | 189 |
| 7 | KPNJ2_RS09410 | conjugal transfer<br>protein TraD | 1,892,386..1,892,709 | 324 | 107 |
| 8 | KPNJ2_RS09405 | hypothetical protein | 1,891,690..1,892,286 | 597 | 198 |
| 9 | KPNJ2_RS27870 | hypothetical protein | 1,891,406..1,891,720 | 315 | 104 |
| 10 | KPNJ2_RS09400 | hypothetical protein | 1,889,929..1,890,411 | 483 | 160 |
| 11 | KPNJ2_RS09395 | MobQ family<br>relaxase | c1,887,562..1,889,082 | 1521 | 506 |
| 12 | KPNJ2_RS09390 | hypothetical protein | c1,887,253..1,887,468 | 216 | 71 |
| 13 | KPNJ2_RS09385 | hypothetical protein | 1,886,838..1,887,080 | 243 | 80 |
| 14 | KPNJ2_RS09380 | tetratricopeptide<br>repeat protein | c1,885,210..1,886,475 | 1266 | 421 |
| 15 | KPNJ2_RS09375 | TIR domain-containing protein | c1,884,789..1,885,208 | 420 | 139 |
| 16 | KPNJ2_RS09370 | hypothetical protein | 1,884,469..1,884,744 | 276 | 91 |
| 17 | KPNJ2_RS09365 | hypothetical protein | 1,883,908..1,884,459 | 552 | 183 |
| 18 | KPNJ2_RS09360 | hypothetical protein | 1,883,327..1,883,893 | 567 | 188 |
| 19 | KPNJ2_RS29065 | hypothetical protein | 1,882,543..1,882,821 | 279 | 92 |
| 20 | KPNJ2_RS09355 | hypothetical protein | c1,880,900..1,881,607 | 708 | 235 |
| 21 | KPNJ2_RS27855 | DUF4756 family<br>protein | c1,880,333..1,880,812 | 480 | 159 |
| 22 | KPNJ2_RS09350 | H-NS family<br>nucleoid-associated<br>regulatory protein | c1,879,770..1,880,174 | 405 | 134 |
| 23 | KPNJ2_RS09345 | hypothetical protein | 1,879,264..1,879,644 | 381 | 126 |
| 24 | KPNJ2_RS09340 | hypothetical protein | 1,878,921..1,879,211 | 291 | 96 |
| 25 | KPNJ2_RS09335 | hypothetical protein | 1,878,531..1,878,842 | 312 | 103 |
| 26 | KPNJ2_RS09330 | hypothetical protein | 1,878,188..1,878,517 | 330 | 109 |
| 27 | KPNJ2_RS09325 | AlpA family<br>transcriptional<br>regulator | 1,877,954..1,878,172 | 219 | 72 |
| 28 | KPNJ2_RS09320 | DUF6387 family<br>protein | 1,876,943..1,877,806 | 864 | 287 |

<sup>a</sup>A BLASTp search identifies the product of this gene as SpnT (WP\_242498927.1;
98% coverage, 99.8% identity).

As we previously reported, a nested PCR approach followed by DNA sequencing showed that ICEKp258.2 can be excised from the chromosome of *K. pneumoniae* strain KP35, reconstituting its integration site (*attB*) at *asn1D*; however, it was not possible to demonstrate circularization of the GI since the putative *attP* site on the excised GI could not be amplified (19). Therefore, we applied a modified approach in which high-fidelity PCR was carried out in two separate and successive rounds but using the same primer pairs, being the template for the second round, a sample from the first one. Following this approach, we were able to obtain amplicons for both *attB* and *attP* after the second PCR round (**Fig. 1C**). The faint bands, or their absence, after the first round indicates that formation of the *attB* and *attP* sites is a low-frequency event. Although two noticeable bands were obtained for *attP*, DNA sequencing demonstrated that only the 667 bp band (the expected size) contained both ends of the GI, joined by an exact copy of the *attB* site, which is only possible as result of the circularization of ICEKp258.2 (**Fig. 1D**). The other PCR product, of higher size, was an unspecific amplification and, as revealed by DNA sequencing, corresponded to a ≈725 bp region spanning the 3’ and 5’ ends of the *rhiT* and *rhiN* genes, located at ≈53.7 kb upstream from ICEKp258.2 (Data not shown).

Our data show that ICEKp258.2 carries a conserved set of genes shared among the EARL GIs, mainly those predicted to encode proteins related to the core functions of excision/integration and transfer. Accordingly, the excision of this GI results in the formation of a circular element, which may potentially be the subject of horizontal transfer.

### Genomic islands closely related to ICEKp258.2 are present among different *K. pneumoniae* sequence types and other species of pathogenic bacteria

The relatively recent acquisition of ICEKp258.2 by ST258 (7) and its excision capacity underscore the transfer potential of this GI. However, it has not been reported outside ST258 and ST512 strains. However, as exemplified in **Fig. 1A**, genomes from different STs harbor GIs with high nucleotide identity relative to ICEKp258.2 (ICEKp258.2-like GIs). We used BLAST to query the non-redundant nucleotide database to identify other *K. pneumoniae* STs or other bacterial species harboring ICEKp258.2 or closely related GIs.

We found 123 genomes harboring 125 GIs with ≥70% alignment coverage to ICEKp258.2 and a nucleotide identity ≥88.2% (**Fig. 2; Supplemental Table S2**). The identified GIs were mainly carried by *K. pneumoniae* strains (107/125), followed by *E. coli* (4/125), *K. grimontii* (3/125), *Citrobacter* sp. (3/125), *Enterobacter* sp. (3/125), *K. quasipneumoniae* (2/125), *Serratia marcescens* (2/125), and *Raoultella ornithinolytica* (1/125) strains. Most *K. pneumoniae* strains belonged to ST258 (56/107), followed by ST11 (12/107) and ST512 (11/107). The other STs had three or fewer representatives.

**Figure 2.**
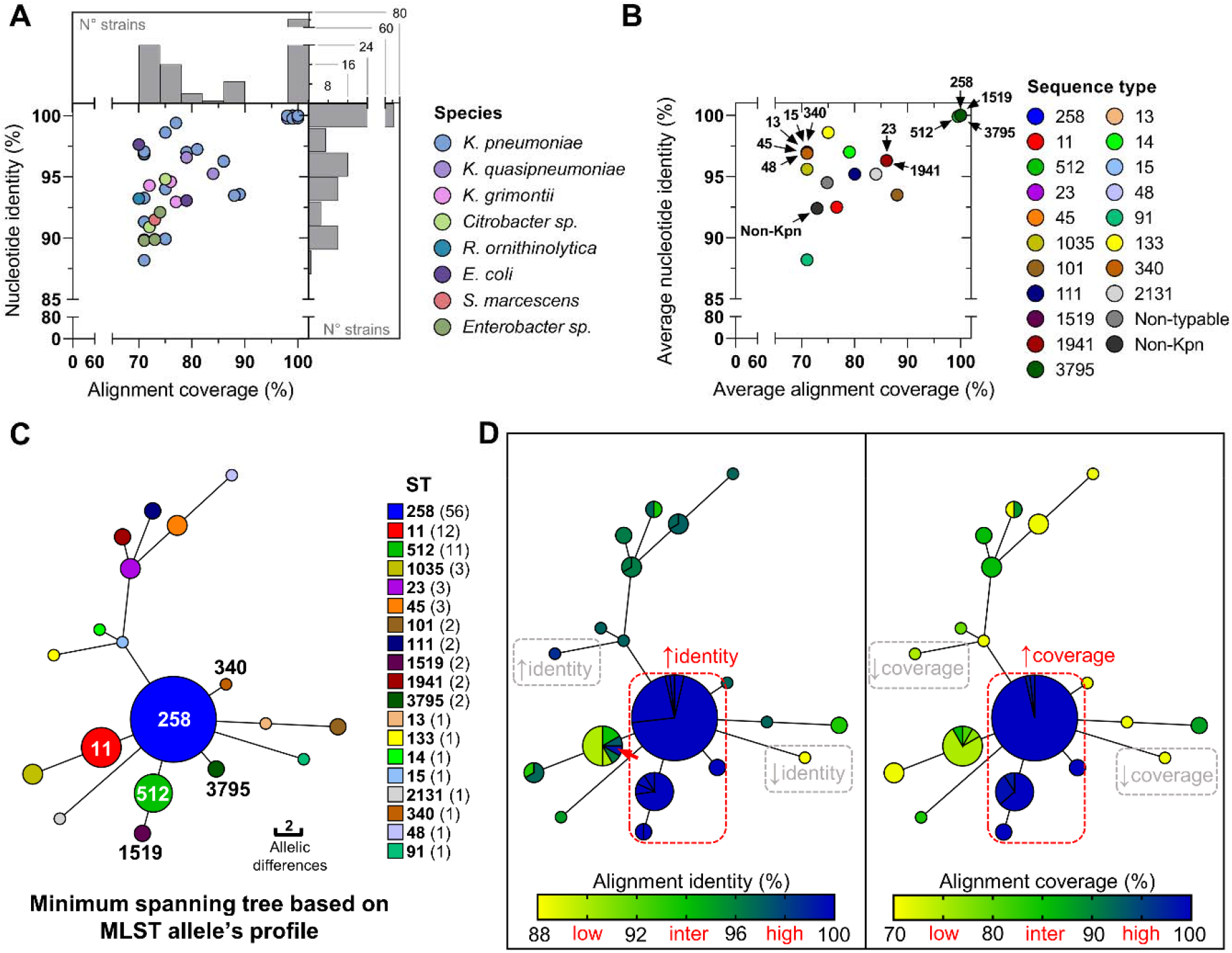
Genomic islands closely related to ICEKp258.2 are present among different *K. pneumoniae* STs and other species of pathogenic bacteria. A) Nucleotide sequence identity and coverage percentage of the alignment between ICEKp258.2 from NJST258_2 and the most closely related GIs in the BLAST non-redundant database. A 70% coverage threshold was used to filter the BLAST results. Colors represent the different bacterial species harboring the identified GIs. Histograms on the right and above the scatter plot show the number of strains harboring a GI with a given nucleotide identity or alignment coverage relative to ICEKp258.2. **B)** Nucleotide sequence identity and coverage percentage but averaged within a given *K. pneumoniae* ST. The colors represent the different STs. Non-typable *K. pneumoniae* isolates or non-*Kp* isolates are indicated in gray and black colors, respectively. **C)** Minimum spanning tree (MST) of the typable *K. pneumoniae* genomes shown in **A** based on the 7-gene MLST. The node size is proportional to the number of genomes included in each ST, indicated in parenthesis on the legend. The edges represent the number of allelic differences between two connected nodes. **D)** The same MST in **C** but colored by the nucleotide identity (left) and coverage (right) of the alignments between the ICEKp258.2 GI from strain NJST258.2 and the closely related GIs harbored by typable *K. pneumoniae*. The red dashed line indicates genomes harboring GIs with both high identity (≥99.4%) and high alignment coverage (≥98%). Examples of genomes carrying GIs with high identity/low coverage, and with low identity/low coverage are also indicated with gray dashed lines.

The majority of the GIs (71/125) had a high alignment coverage (≥98%) and sequence identity (≥99.8%) with ICEKp258.2 and were carried by *K. pneumoniae*, therefore, were considered as the same GI (**Fig. 2B**). All genomes from ST258 and ST512, an SLV of ST258, carried ICEKp258.2. Interestingly, we also found this island in genomes from ST3795 and ST1519, which are SLV and double locus variants (DLV) of ST258 (**Fig. 2B, 2C**). No other species, genus, or ST were found to harbor ICEKp258.2. The GIs with the second highest sequence identity with ICEKp258.2 (96.3%-99.4%) were found mainly in *K. pneumoniae* genomes from a diversity of STs (e.g., 11, 14, 15, 23, 45, 48, 133, 340, among others), and in two *K. quasipneumoniae* and one *E. coli* genomes (**Supplemental Table S2**). However, despite the high sequence identity, the alignment coverage of the GIs from this second group was low and ranged from 70% to 81%. The GIs with an intermediate alignment coverage (81%-89%) had high nucleotide identities ranging from 93.4% to 97.2% and were carried by *K. pneumoniae* genomes from STs 11, 23, 101, 111, 1941, and 2131 (**Fig. 2D**). These GIs with high/intermediate alignment coverage and high nucleotide identity are different from ICEKp258.2 and could represent ancestors or descendants of this island. Noteworthy, an ST11 strain harbored the GI with the highest nucleotide identity with ICEKp258.2, the same ST that originated the ancestral ST258 (**Fig. 2D; Supplemental Table S2**).

### ICEKp258.2 likely derived from a genomic island carried by an ST11 *K. pneumoniae* strain

To assess the phylogenetic relationships between ICEKp258.2 and the identified GIs, we built a maximum likelihood tree based on the concatenated nucleotide sequences of the integrase-and RDF-encoding genes. The 125 GIs were filtered to exclude duplicated GIs. However, if the same GI was present in a different genus, species, or ST, it was included in the analysis.

Overall, ICEKp258.2 located near the base of the tree, having as closest relatives the GIs harbored by other *K. pneumoniae* strains from STs 11, 14, 45 y 133, which are the same GIs that had the highest overall nucleotide identity (≥96.6%) with ICEKp258.2 (**Fig. 3A**). As moving away from the base of the tree, the diversity of host bacteria increases, as expected from the horizontal dissemination of these GIs. A putative recent horizontal gene transfer event could be evidenced between strains F16KP0075 (ST11), KP7 (ST1941), and ED2 (ST23), as these bacteria carry the same GI, which also shares a high nucleotide identity (96.3%) and intermediate alignment coverage (86%) with ICEKp258.2. Another putative transfer event, this time between different genera, could also be evidenced between strains *K. pneumoniae* BeachRanger (ST11) and *Enterobacter roggenkampii* OIPH-N260. All GIs carried the genes encoding the type-III restriction-modification system and T4P-related proteins. The genes encoding the putative spnT (*spnT*) protein and the H-NS (*hfp*) homolog were present in all GIs except two for *spnT* and four for *hfp*. Genes encoding putative TIR-domain-containing proteins could be identified in six GIs besides ICEKp258.2.

**Figure 3.**
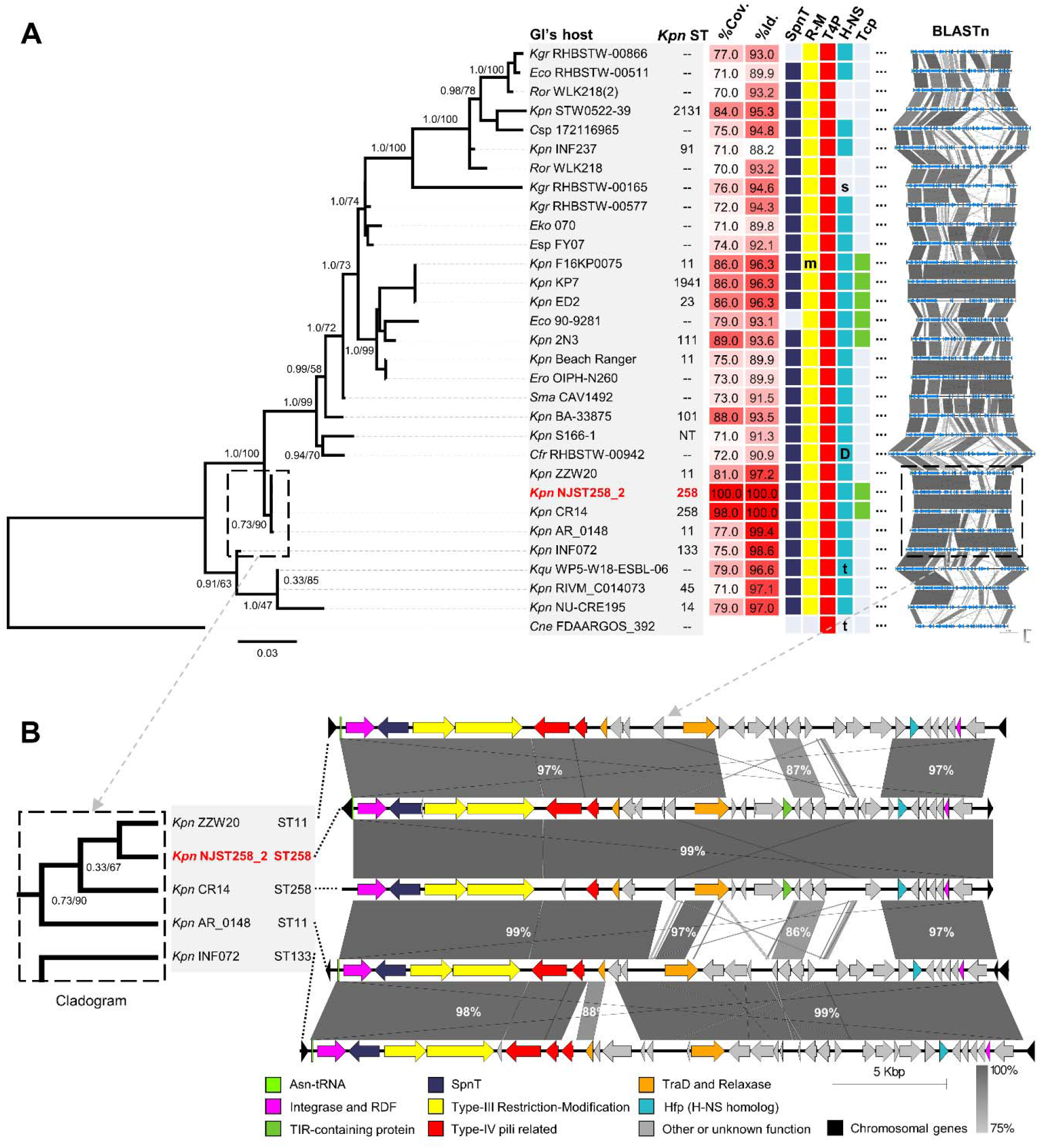
ST258 acquired ICEKp258.2 likely from a ST11 strain. A) Maximum likelihood phylogenetic tree of ICEKp258.2 and its closely related GIs based on the alignment of the concatenated nucleotide sequence of the integrase and RDF encoding genes The corresponding sequences of the EARL GI from *Cedecea neteri* strain FDAARGOS_392 were used as an outgroup. The numbers near the nodes indicate the approximate Bayes and Ultrafast bootstrap (5000 replicates) support only for basal nodes. From left to right, the panel also shows: the ST (only for typable *K. pneumoniae*); the nucleotide identity and coverage from the alignment of each island with ICEKp258.2; the presence of the genes predicted to encode SpnT, the Type-III restriction modification system (R-M; the m indicates a disruption of the methylase-encoding gene), the type-IV-pili-related proteins (T4P), the H-NS homologs (full-length, colored square; short, s; truncated, t; or duplicated, D) and the TIR-domain-containing proteins; and the nucleotide identity (BLASTn) across the analyzed GIs. **B)** A cladogram view of the branch including the ICEKp258.2 GI and its closely related GIs, and the corresponding BLASTn alignments of the entire islands. The percentages indicate the nucleotide identity between the island regions.

According to the phylogeny, the closest relatives to ICEKp258.2 were the GIs carried by strains ZZW20 and AR_0148, both from ST11 (**Fig. 3A**). A cladogram representation of this group evidenced that the GI from the strain AR_0148 was basal to ICEKp258.2 (**Fig. 3B**), which is the result of only two nucleotide differences in the RDF-encoding gene at positions 78 and 138 of the coding sequence. A more detailed examination of the sequences of the GIs carried by strains CR14, NJST258_2, ZZW20, and AR_0148 was carried out. The BLASTn alignments (**Fig. 3B**) of the complete GIs revealed a 99% nucleotide identity between one half of the GI from AR_0148 and ICEKp258.2. Two other regions, the relaxase-encoding gene and the last eight genes from the GI carried by AR_0148, shared 97% nucleotide identity with ICEKp258.2. Except for three genes with 86% identity, the remaining sequence of the GI in AR_0148 had a nucleotide identity below 75%. Besides the islands harbored by the ST11 strains ZZW20 and AR_0148, the GI carried by strain INF072 (ST133) had the highest nucleotide identity with ICEKp258.2 (98.6%), although limited to 75% of the GI sequence. Noteworthy, this GI from strain INF072 included almost the entire sequence of the AR_0148 GI (**Fig. 3B**). These results support the idea of ICEKp258.2 as being acquired from an ST11 strain. However, the sequence alignment between pairs of the ICEKp258.2-like GIs (see BLASTn in **Fig. 3**, **Supplemental Fig. S1**) shows an interspersed distribution of high and low identity regions, which is indicative of modular recombination events between similar GIs, as those low identity regions are present with high identity in other islands. Such modular recombination may obscure the phylogenetic signal.

### Subinhibitory concentrations of clinically relevant antibiotics promote the excision of ICEKp258.2 and modulate gene expression in the island

The excision of GIs can lead to their horizontal transfer and also be implicated in regulating gene expression within the islands (22–24). Therefore, we aimed to characterize the excision of ICEKp258.2 further using the GI carried by *K. pneumoniae* ST258 strain KP35 as a model. First, we estimated the proportion of the bacterial population with an excised GI in common culture conditions, i.e., LB medium at 37°C (**Fig. 4A**, 4B). We found that the excision of ICEKp258.2 is an event of very low frequency, in concordance with the need for two rounds of PCR to obtain visible bands of the *attB* and *attP* regions (**Fig. 1C**). At the logarithmic (log) phase, an average 4×10^-5^% of the population (40 out of 10^6^ bacteria) had an excised ICEKp258.2. The proportion increased 2.5X in the stationary phase (10^-4^%; or 100 out of 10^6^ bacteria) (**Fig. 4C**). This increase in the excision was accompanied by a downregulation of the expression within the island by 2-7 log_2_ (**Fig. 4D**), indicating that the higher the excision, the lower the gene expression. Correlation analysis and linear regression showed that all the correlation coefficients (Pearson r) were negative and that for 4 out of 6 genes (*int*, *spnT*, *res*, and *tcp*), the slope significantly deviated from zero (**Fig. 4E**).

**Figure 4.**
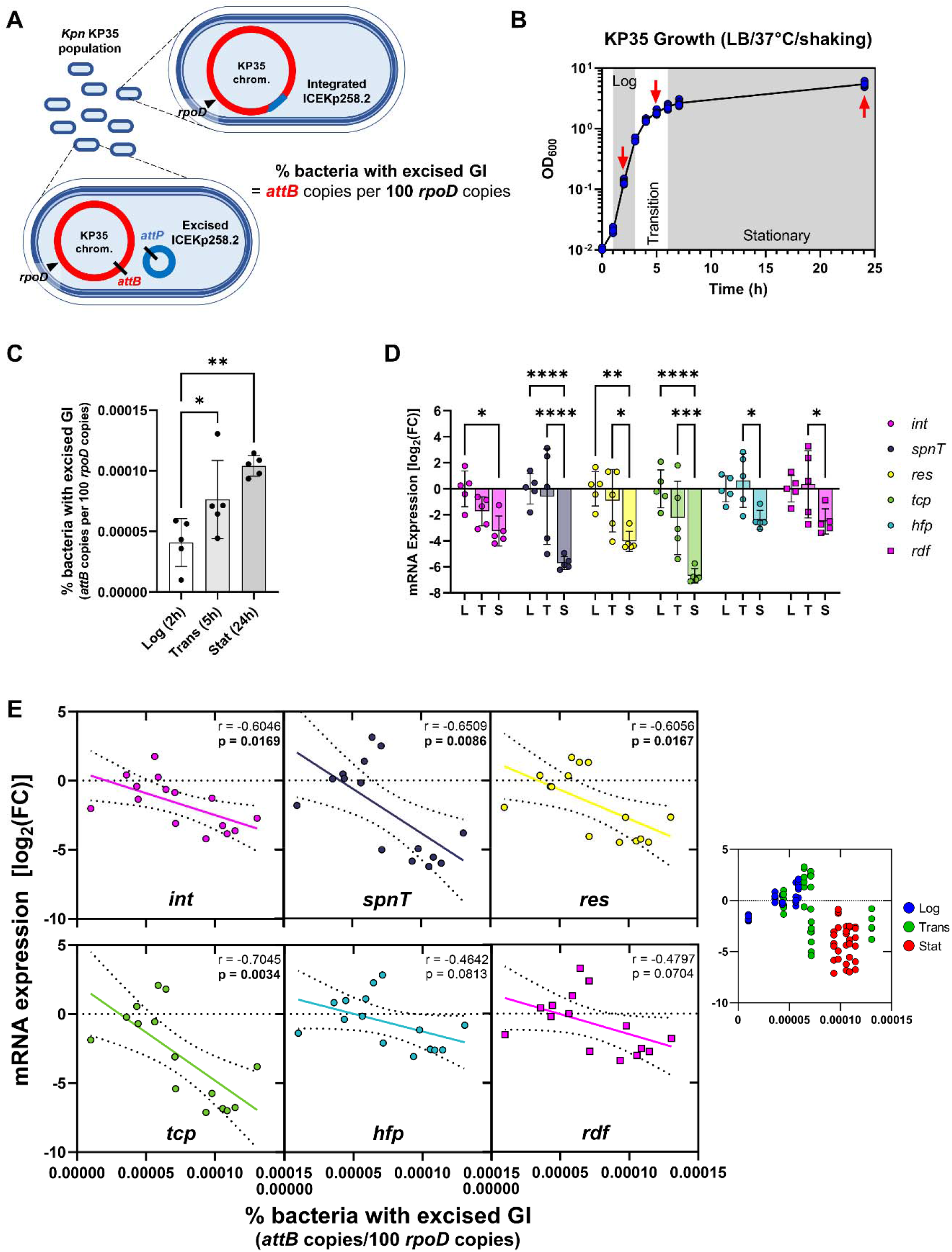
The expression within ICEKp258.2 negatively correlates with the excision of the island. A) Schematic representation of the integrated and excised states of ICEKp258.2 within a population of *K. pneumoniae* ST258 strain KP35. Quantification of the *attB* copies per 100 *rpoD* copies approximates the percentage of bacteria with an excised GI. **B)** Growth curves (5 biological replicates) of strain KP35 in LB. The red arrows indicate the times in which the samples were collected for genomic DNA and total RNA extraction. **C)** Percent KP35 with an excised ICEKp258.2 at the different phases of growth in LB. **D)** mRNA expression of selected ICEKp258.2 genes relative to the expression at the log phase of growth in LB. **E)** Scatter plots of the expression levels of selected genes within ICEKp258.2 versus the excision of the GI. Continuous lines represent the linear regression of the data. The seventh scatterplot at the right shows all the data points colored according to the growth phase. Statistical significance for excision data (C) was assessed through one-way ANOVA followed by Dunnett’s multiple comparisons test. For expression data (D), two-way ANOVA was used, followed by Tukey’s multiple comparison tests. Scatterplots show the Pearson correlation coefficient and the p-value. *p<0.05, **p<0.01, ***p<0.001, ****p<0.0001.

In the environment and host tissues, bacteria are exposed to subinhibitory concentrations of antibiotics due to agricultural activities, wastewater treatment, or incorrect therapy in the clinical setting, among other reasons (25). This phenomenon has been shown to impact virulence expression and the interaction with the host immune system in different pathogens, including *K. pneumoniae* (26–28). Moreover, the horizontal transfer of some GI families is promoted in the presence of antibiotics by the activation of the excision and transfer machinery (29–31). Therefore, we assessed the impact of subinhibitory concentrations of antibiotics relevant in the treatment of *K. pneumoniae* infections that also targeted different cellular processes: Cell wall synthesis (ceftazidime/avibactam, CZA; meropenem, MEM; imipenem, IPM); protein synthesis (chloramphenicol, CHL; gentamicin, GEN); and DNA replication (ciprofloxacin; CIP) (**Fig. 5A**, **Supplemental Fig. S2**).

**Figure 5.**
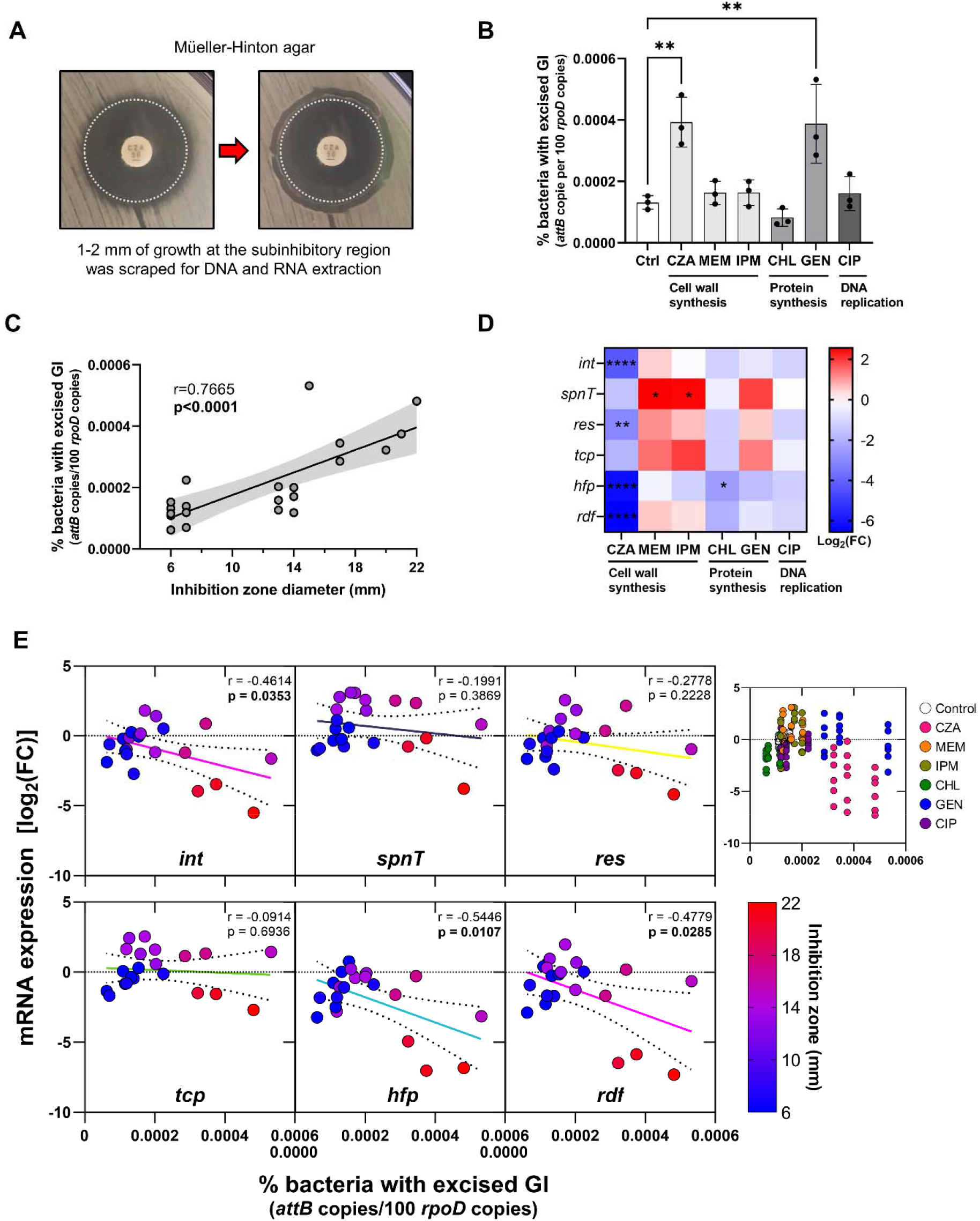
Antibiotic potency positively correlates with the excision of ICEKp258.2. A) Photograph showing how the bacterial growth occurring next to the border of the antibiotic inhibition zone (subinhibitory region) was scraped to obtain the samples for genomic DNA and total RNA extraction. Three biological replicates were carried out. The dashed white circle indicates the limit of the inhibition zone. **B)** Percent KP35 with an excised ICEKp258.2 at different subinhibitory antibiotic concentrations on Müeller-Hinton agar. The antibiotics and their cell target are indicated in the X axis (Ctrl: control; CZA: ceftazidime-avibactam; MEM: meropenem; IPM: imipenem; CHL: chloramphenicol; GEN: gentamycin; CIP: ciprofloxacin). **C)** Scatter plot showing the positive correlation between the diameter of the inhibition zone and the excision of ICEKp258.2. The black line represents the linear regression of the data, and the light gray area indicates the 95% confidence bands. **D)** Heatmap showing the mRNA expression of selected ICEKp258.2 genes, under subinhibitory antibiotic concentrations, relative to the expression in the absence of antibiotics. **E)** Scatter plots of the expression levels of selected genes within ICEKp258.2 versus the excision of the GI. Continuous lines represent the linear regression of the data. The seventh scatterplot at the right shows all the data points colored according to the antibiotic treatment. Statistical significance for excision data (B) was assessed through one-way ANOVA followed by Dunnett’s multiple comparisons test. For expression data (D), two-way ANOVA was used, followed by Dunnett’s multiple comparison tests. Scatterplots show the Pearson correlation coefficient and the p-value. *p<0.05, **p<0.01, ***p<0.001, ****p<0.0001.

The proportion of bacteria with an excised ICEKp258.2 after 18-20h of incubation at 37°C in Mueller-Hinton II agar was similar to that observed in LB after 24h (1.3×10^-4^% in MH and 10^-4^% in LB; Ctrl in **Fig. 5B**). Interestingly, the proportion of KP35 with an excised ICEKp258.2 increased 3X when challenged with subinhibitory concentrations of CZA and GEN (3.9×10^-4^% and 3.8×10^-4^%, respectively; **Fig. 5B**). On the other hand, MEM, IPM and CIP induced a slight increase, and CHL a slight decrease in excision that was not statistically significant. Noteworthy, while there was no correlation between the excision level and the antibiotic target or the antibiotic class (**Fig. 5B**), there was a positive correlation (r=0.77) between the excision and the antibiotic potency, expressed as the diameter of the inhibition zone (**Fig. 5C**).

The expression within ICEKp258.2 was altered by the antibiotics at subinhibitory concentrations in a gene-and antibiotic-dependent manner (**Fig. 5D**, **Supplemental Fig. S3A**). The genes *int*, *res*, *hfp,* and *rdf* were downregulated (≤-3.1 log_2_) in the presence of CZA, while CHL caused a decrease in the expression of *hfp* only (≤-2.5 log_2_). On the other hand, MEM and IPM increased the expression of *spnT* (≥2.5 log_2_). The effect of the other antibiotics was not statistically significant, although a trend was observable for MEM, IPM, and GEN to cause a slight upregulation of *spnT*, *res,* and *tcp* (0.5-1.9 log_2_). While a higher excision correlated with a lower gene expression in the absence of antibiotics in LB (**Fig. 4E**), the expression level in the presence of subinhibitory concentrations of the tested antibiotics was more dependent on the gene as neither the excision nor the antibiotic potency consistently correlated with gene expression, and only the *int*, *hfp,* and *rdf* genes invariably had a negative correlation between their expression and the excision or potency (**Fig. 5E**, **Supplemental Fig. S3B**). These results show that antibiotics relevant to the treatment of *K. pneumoniae* infections can modulate the excision of ICEKp258.2 and the gene expression within this GI.

## DISCUSSION

Carbapenem-resistant *K. pneumoniae* ST258 is a highly successful multidrug-resistant lineage associated with increased treatment costs and poor treatment outcomes. However, the factors involved in its emergence and widespread dissemination remain poorly understood (10, 11). ICEKp258.2 is an excisable genomic island hypothesized to play a relevant but unknown role in the success of ST258 as it was acquired through horizontal gene transfer by the ancestral ST258 strain from an unknown donor before global dissemination, and as it carries genes encoding putative factors involved in adherence to surfaces and in shaping the plasmid pool of ST258 (10, 12, 13). Here, we presented a first characterization of ICEKp258.2, analyzing its gene content, the GI distribution and phylogenetic relationships, and the GI response to clinically relevant antibiotics.

We found that the ICEKp258.2 gene content is shared by other members of the Enterobacterales-associated ROD21-like family of GIs, especially the excision/integration and putative conjugation modules, the T4P-related genes and, to a lesser extent, the type-III RM genes. Conservation of the genes from the excision/integration module (integrase and RDF) and conjugation module (TraD and MOB_Q_ relaxase) are characteristic of the Enterobacterales-associated ROD21-like family of GIs (19, 32), as expected for the genes encoding the major functions related to a GI’s biology: excision/integration and transfer, which mediate dissemination. This is in agreement with the high diversity of bacterial families and species in which the EARL GIs, including ICEKp258.2, are found (19). Different authors have suggested that the T4P-related genes and those from the type-III RM system in ICEKp258.2 could play key roles in the success of ST258. We previously found that the T4P-related genes are also characteristic of the EARL GIs but not as conserved as the excision/integration and transfer modules (19), suggesting a beneficial role for the host instead of directly for the GI itself. In the foodborne pathogen *Salmonella enterica* serovar Enteritidis the T4P-related genes are carried by ROD21, the most studied member of the EARL family (33), and are required for the successful colonization of murine internal organs as their deletion results in a reduced bacterial load in the liver and spleen (34). No other studies have addressed the role of the T4P-related genes of the EARL GIs. Less is known about the role of the ICEKp258.2-encoded type-III RM system. However, we previously found that the acquisition of the type-III RM genes by the EARL GIs coincided with their presence in a higher diversity of bacterial hosts. While the EARL GIs lacking the type-III RM genes are carried by *Pectobacterium* spp., *Escherichia coli,* and *Salmonella* serovars, the type-III RM positive GIs are found in *E. coli*, *Salmonella*, *Citrobacter*, *Enterobacter*, *Raoultella*, *Serratia*, *Yersinia*, *Cedecea*, *Kluyvera* and *Klebsiella* strains (19). A similar diversity of hosts harbored the ICEKp258.2-like GIs identified in this study, and all carried the type-III RM genes. These findings suggest a role of the type-III RM system in the dissemination of the EARL GIs and possibly for ICEKp258.2. In this context, two hypotheses arise: i) as RM systems can restrict the acquisition of foreign genetic elements by recognizing and cleaving non-methylated DNA, the RM-positive EARL GIs could restrict the pool of horizontally acquired elements to favor the presence of plasmids (or an unknown genetic element) that mediate their conjugative transfer; and ii) the RM-positive EARL GIs produce genetic addiction on the bacterial host by post-segregational killing that favor survival of the bacterial cells that successfully integrate the GI and therefore methylate their DNA to avoid cleavage by the restriction endonuclease (35, 36). The high prevalence of the T4P-related and type-III RM genes among the EARL GIs highlights them as suitable targets for deletion experiments aiming to study their role in the success of ST258. While the multidrug-resistant profile of ST258 strains severely limits the tools to perform genetic engineering in these pathogens, recent techniques coupling unusual antibiotics such as Zeocin, the lambda RED system, and CRISPR-Cas9 have been successfully applied to perform deletions and other modifications in ST258 (11, 37).

We report here that ICEKp258.2 is not confined to ST258 and its SLV ST512 but is also present in strains from ST3795 and ST1519, *rpoB222,* and *gapA54*/*rpoB9* variants of ST258, respectively. Noteworthy, recent reports from Italy show that ST1519 includes multidrug-resistant strains associated with ceftazidime/avibactam (CZA) resistance and with *bla_KPC_*variants differing from the *bla_KPC-2_* and *bla_KPC-3_* variants usually associated with ST258 (38–41). Acquisition of ICEKp258.2 by these isolates likely resulted from vertical transmission, as phylogenetic and comparative analyses indicate that ST1519 stems from ST258 (39). Besides ST258 and its single and double locus variants, ICEKp258.2 was not found in any other ST. However, we found many GIs sharing high nucleotide identity among other *K. pneumoniae* STs, other *Klebsiella* species, and other genera. The rooted phylogeny, based on the concatenated integrase and RDF gene sequences, unveiled events of horizontal gene transfer of these GIs between strains from different STs (ST11, ST23, ST1941) and even between different genera (*K. pneumoniae* ST11 and *Enterobacter roggenkampii*). Importantly, according to the metadata associated with the genome accession number or published literature, the strains mentioned above include clinical multidrug-resistant isolates from invasive infections (42, 43). Additionally, the phylogeny of the ICEKp258.2-like GIs placed the GIs from strains ZZW20 and AR_0148 as the closest relatives to ICEKp258.2, being also the GIs with the highest overall nucleotide identity (97.2% and 99.4%), and the GI harbored by AR_0148 being basal in the clade. However, the evidence for modular recombination events between the different GIs indicates that the reconstructed phylogeny must be interpreted cautiously. Nevertheless, as ZZW20 and AR_0148 are from the same ST that originated ST258, our findings suggest that ST258 acquired ICEKp258.2 from an ST11 strain. Our analyses also suggest active transfer of the ICEKp258.2-like GIs (also members of the EARL GIs) between bacterial pathogens of clinical relevance.

The excision of ICEKp258.2 increased during the transition and stationary phases of growth. Similar results were previously found for the EARL GI ROD21 and many other GIs from different island families (31, 32, 44). While there is evidence indicating that for some GIs, such as ICE*Ml*Sym^R7A^ from *Mesorhizobium loti* R7A, quorum sensing can play a role in modulating excision (44), for most GIs, the mechanistic details for this phenomenon are not clear. To date, there is no evidence that ICEKp258.2 or other EARL GIs encode proteins related to quorum sensing. Different environmental stimuli can also promote the excision and transfer of GIs, including pH, temperature, host factors, UV radiation, and antibiotics (23, 29, 45, 46). Antibiotics with DNA-damaging capacity, such as mitomycin C, ciprofloxacin, metronidazole, and others, can activate the SOS response leading to LexA-mediated cleavage of GI-encoded repressors, thus inducing the excision machinery of the islands (30, 31, 47). However, the excision of ICEKp258.2 was not induced by ciprofloxacin but by the cell wall-synthesis inhibitor ceftazidime/avibactam (CZA) and the protein-synthesis inhibitor gentamicin (GEN) at subinhibitory concentrations. The observed induction seems unrelated to the targeted pathway as the other tested antibiotics, also targeting cell wall and protein synthesis, had no effect on excision. Importantly, the positive correlation between antibiotic potency and the excision level suggests that the increased ICEKp258.2 excision in the presence of CZA and GEN, which produced the larger inhibition zones, could be an indirect result of the antibiotic-induced stress response that includes the release of reactive oxygen species (48). The exposure to subinhibitory antibiotic concentrations also resulted in altered gene expression within ICEKp258.2 without a clear trend that could relate the expression changes to antibiotic class or targeted pathway. Nonetheless, our analysis evidenced that gene expression responses were dependent on the gene/antibiotic combination. A limitation of our study is that we cannot link the observed changes in excision or gene expression with phenotypical alterations that could impact the virulence of *K. pneumoniae*. Efforts for implementing the CRISPR-Cas9-based editing tool for the study of ICEKp258.2, and other GIs, are currently being carried out in our laboratory.

The emergence of bacterial pathogens is a multifactorial and complex phenomenon in which environmental, host and bacterial factors interact and participate in the development/acquisition of virulence traits. This process is complemented by the emergence of antibiotic resistance (49). Identifying the factors involved in pathogen emergence has major public health implications as their study can contribute to develop surveillance strategies and control measures (10, 49). Our study provides the basis to understand the biology of ICEKp258.2, placing it within the EARL family of GIs and uncovering a potential role of antibiotics in the spread of the ICEKp258.2-like GIs among bacterial pathogens.

## MATERIALS AND METHODS

### Sequences of ICEKp258.2 and the EARL GIs and assessment of gene conservation

The sequence of ICEKp258.2 and the other 55 EARL GIs previously identified were downloaded from GenBank according to the accession numbers and genomic coordinates previously published (19). The sequence of ICEKp258.2 from *K. pneumoniae* ST258 strain NJST258_2 (GenBank accession NZ_CP006918.1; nucleotides 1876030 to 1904524) was selected to perform the analyses. The downloaded sequences included the entire GIs with additional 500 bp upstream and downstream of the islands. The EARL GIs found in unannotated genomes were downloaded and annotated using RAST at https://rast.nmpdr.org/rast.cgi. Global sequence alignments and tBLASTx alignments were performed with ICEKp258.2 and the EARL GIs using Mauve v20150226 and EasyFig v2.2.2. The proportion of the ICEKp258.2 genes among the other EARL GIs was represented as a percentage of the EARL GIs carrying a homolog of the corresponding gene (Table S1).

### Identification of ICEKp258.2-related GIs, distribution, and phylogenetic analysis

To search and identify GIs closely related to ICEKp258.2, the downloaded sequence of ICEKp258.2 was used to perform a BLASTn search against the non-redundant nucleotide database. Hits with ≥70% sequence coverage and ≥88.19% sequence identity were selected for further analysis. The boundaries of each GI were manually examined for the presence of the characteristic genes located at the distal regions of the EARL GIs: a P4-related integrase downstream of the insertion site, and a putative RDF followed by a conserved gene of unknown function on the other side. The bacterial host species, strain name, genome accession number, alignment coverage, alignment identity, location within its host genome, and comments regarding the insertion site (*asn1A*, *1B*, *1C,* or *1D*) and the presence of insertion sequences within the island were recorded (Table S2). For GIs harbored in *K. pneumoniae*, the sequence type of their hosts was assigned using PubMLST (https://pubmlst.org/).

A minimum spanning tree (MST) was constructed for the *K. pneumoniae* genomes harboring ICEKp258.2 or related GIs using GrapeTree and the MSTree V2 algorithm (50), based on the allelic profile of the 7-gene MLST scheme. The same MST was colored according to sequence type, and percent coverage and nucleotide identity of the alignment between the harbored GI and ICEKp258.2.

For the phylogenetic analysis, the GIs identified by BLASTn were manually filtered to exclude duplicated GIs (the same GI in a different genome). Putative duplicated GIs were first identified based on highly identical percent coverage and identity, the nucleotide length, and presence in the same bacterial species. Then, BLAST was used to confirm that they were the same GI. For selected GIs, the nucleotide sequences from the integrase-and RDF-encoding genes were concatenated, and a multiple codon-based nucleotide alignment was obtained using MUSCLE in MEGAX. The corresponding genes from the EARL GI carried by *Cedecea neteri* FDAARGOS_392 (GenBank accession CP023525.1) were included in the alignment and used as an outgroup. The best-fitting nucleotide substitution model was selected for each partition based on the Bayesian Information Criterion (TIM3e+I+G4 and TVM+F+I), and a maximum likelihood tree was constructed using the IQ-TREE web server (http://iqtree.cibiv.univie.ac.at/). The node support was calculated by applying the approximate Bayes option and from 5000 bootstraps using the Ultrafast bootstrap option. The aligned sequences from the GIs carried by strains ZZW20, CR14, and NJST258_2 were identical. However, the algorithm of IQ-TREE excluded ZZW20 (for being identical) during tree construction and then added it at the end, creating the artifact of the GI from strain CR14 being basal to those from NJST258_2 and ZZW20 in the cladogram representation. The sequence from the KP7 GI, identical to those from F16KP0065 and ED2, was subjected to the same treatment by IQ-TREE. The figure of the phylogenetic tree was made using FigTree v1.4.3.

### Bacterial strains, culture media, and antimicrobial disks

*K. pneumoniae* strain KP35 and strain KPPR1 were selected as a carbapenem-resistant ST258 model and an antibiotic-susceptible control, respectively (17). Both strains were maintained at-80°C in CRYOBANK vials. When required, bacteria were grown overnight in LB at 37°C with shaking. Exposure to subinhibitory antimicrobial concentration was carried out in Mueller-Hinton II agar (Becton-Dickinson). Antimicrobial disks for ceftazidime/avibactam (CZA; 30/20 µg), imipenem (IMP; 10 µg), chloramphenicol (CHL; 30 µg), meropenem (MEM; 10 µg), gentamycin (GEN; 120 µg) and ciprofloxacin (CIP; 5 µg) were purchased and stored at 4-8°C. CZA, IMP, CHL, and MEM were from Oxoid, GEN from Mast, and CIP from Farmalatina.

### Assessment of ICEKp258.2 circularization after excision

An overnight culture of *K. pneumoniae* KP35 was prepared, and 1 mL was centrifuged at 8000×g for 6 minutes at 4°C. Genomic DNA was extracted, purified, and stored as described in the “DNA and RNA extraction” section (see below). Primers Kpn-2_Fw + Kpn-4_Rev (519 bp) and Kpn-4_Fw + Kpn-2_Rev (667 bp), and the Platinum SuperFi II high-fidelity DNA polymerase (Invitrogen) were used to amplify the *attB* and *attP* regions by PCR. Two rounds of PCR were carried out. For the first round, 50 ng of genomic DNA was used as a template. For the second round, 1 µL of the reaction mix from the first round was used as a source of templates. Each PCR round comprised 35 amplification cycles and used the same primers and polymerase. The amplification products were visualized by electrophoresis at 90V for 55 min in TAE buffer, using 1% agarose with 1X SafeView Plus (Fermelo Biotec). The PCR products were purified using the NucleoSpin Gel and PCR clean-up kit (Macherey-Nagel) according to the manufacturer instructions. The purified PCR products were sequenced (Sanger) in the Unidad de Secuenciación y Tecnologías Ómicas from Pontificia Universidad Católica de Chile.

### Assessment of the excision of ICEKp258.2 and gene expression within the island during the growth phases

A 250 mL flask with 50 mL of LB was inoculated with an overnight culture of *K. pneumoniae* KP35 at an initial OD_600_=0.01. The culture was incubated at 37°C with shaking for 24 h, and the OD_600_ was recorded each hour until 7 h and then at 24 h. Two samples of 1.6 mL were taken at 2, 5, and 24 h of incubation and centrifuged at 8000×g for 7 minutes at 4°C. The supernatant was discarded, and the bacterial pellets were stored at-30°C for DNA extraction or resuspended in 1 mL of TRIzol (Invitrogen) and stored at-80°C for RNA extraction. Five independent biological replicates were performed.

### Assessment of the excision of ICEKp258.2 and gene expression within the island under subinhibitory concentrations of different antibiotics

Fourteen-centimeter diameter Petri plates containing Müeller-Hinton II Agar were inoculated with a 0.1_OD600_ suspension of *K. pneumoniae* KP35, prepared from an overnight culture, using a cotton swab. Immediately, the disks containing CZA, IMP, CHL, MEM, GEN, and CIP were placed on the inoculated agar surface, and the plates were incubated at 37°C for 24 h. After incubation, 1-2 mm of the bacterial growth located next to the perimeter of the inhibition zone or next to the perimeter of the disk (in case of no inhibition) was scraped and resuspended in 100 µL of sterile PBS (Fig. S2). Then 50 µL of this suspension was transferred to a different tube. One sample was stored at-30°C, and the other was resuspended in TRIzol and stored at-80°C. Three independent biological replicates were performed. *K. pneumoniae* KPPR1 was used as a susceptible control to assess the potency of the antibiotics tested.

### DNA and RNA extraction

The genomic DNA (gDNA) and total RNA were extracted and purified as previously reported (32). Briefly, the bacterial pellets stored at-30°C, were lysed in 550 µL of Tris-EDTA buffer (pH 8.0) containing approximately 0.2 mg/mL of proteinase K, 0.05 mg/mL of RNase A, and 0.5% SDS, at 37°C for 1 h. The gDNA was extracted with basic phenol:chloroform:isoamyl alcohol (Winkler), precipitated by the addition of 0.1 volume of 3 M sodium acetate (pH 5.2) plus 1 volume of propan-2-ol followed by incubation at-20°C, and collected by centrifugation at 21000×g during 30 min at 4°C. The precipitated DNA was washed with 500 µL of 75% ethanol, resuspended in 50 µL of nuclease-free water and stored at-30°C.

Purification of total RNA from samples in TRIzol stored at-80°C was carried out as recommended by the manufacturer. The RNA was resuspended in 50 µL of nuclease-free water and stored at-80°C until DNase treatment. Residual DNA was eliminated from the RNA samples using the Turbo DNA-free kit (Invitrogen) according to the manufacturer instructions. The DNA-free RNA was stored at-80°C.

### Joining of the ICEKp258.2 *attB* insertion site and the *rpoD* gene from KP35 and preparation of the standard curve for excision quantification

The *rpoD* gene and the *attB* site (left after excision of ICEKp258.2) from *K. pneumoniae* KP35 were amplified by PCR using the Platinum SuperFi II High Fidelity DNA Polymerase (Invitrogen), and then joined by Gibson assembly (Fig. S4). The *rpoD* gene was amplified in a 35-cycle PCR with the primers Kp-rpoD_Gibson_Fw + Kp-rpoD-attB_Gibson_Rev, obtaining a 1865 bp product. The chromosomal region surrounding the *attB* site was amplified by nested PCR with primers Kpn-1_Fw + Kpn-3_Rev (1004 bp) and primers Kp-rpoD-attB_Gibson_Fw + Kp-attB_Gibson_Rev (931 bp). Both products were gel-purified using the NucleoSpin Gel and PCR clean-up system (Macherey-Nagel) and stored in 5 mM Tris-HCl pH 8.5 at-30°C. The purified products were ligated to obtain the *rpoD*-*attB*_KP35 fragment using the NEBuilder HiFi DNA assembly master mix (New England Biolabs). To obtain a high yield, the *rpoD*-*attB*_KP35 fragment was amplified from the assembly reaction by PCR using the Platinum SuperFi II High Fidelity DNA Polymerase and the primers Kp-rpoD_Gibson_Fw + Kp-attB_Gibson_Rev (2776 bp), and then gel-purified using the NucleoSpin Gel and PCR clean-up kit. The purified *rpoD*-*attB*_KP35 fragment was stored in 5 mM Tris-HCl pH 8.5 at-30°C, until use.

Serial 10-fold dilutions ranging from 10^8^ to 10^2^ copies of the *rpoD*-*attB*_KP35 fragment were prepared in nuclease-free water and stored at-30°C in aliquots of 10 µL. Immediately before excision quantification, a set of 10^8^-10^2^ aliquots were thawed, homogenized, and a 10^1^ and 10^0^ dilutions were prepared to be also included in the qPCR assay.

### Quantification of the ICEKp258.2 excision

The percentage of the bacterial population with an excised ICEKp258.2 was calculated by quantifying the number of *attB* copies per 100 *rpoD* copies by qPCR in each sample. qPCR was carried out using the TaqMan Fast Advanced Master Mix (Applied Biosystems), the primers attB1-KpRT_Fw + attB1-KpRT_Rev and the MGB probe attB-1_RT for detection of *attB*; and the primers rpoD-KpRT_Fw + rpoD-KpRT_Rev and the MGB probe rpoD-KpRT_Probe for detection of *rpoD*. Threshold cycles were interpolated in standard curves for both *attB* and *rpoD* prepared with the *rpoD*-*attB*_KP35 fragment ranging from 10^8^ to 10^0^ copies. Standard curves were run in every batch of reactions. qPCR was performed with two technical replicates in each plate in a StepOnePlus Real Time PCR System.

The excision level, expressed as the percentage of the population with an excised ICEKp258.2 was calculated with the following formula:

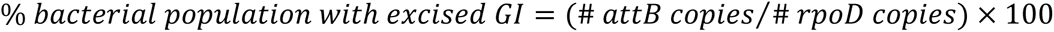

### Quantification of gene expression

The gene expression within ICEKp258.2 was quantified by RT-qPCR for the genes *int*, *spnT*, *res*, *tcp*, *hfp* and *rdf* using the primers Kp-*gene*_RT_Fw/Rev and rpoD-KpRT_Fw/Rev (Table S4). cDNA was synthesized from total RNA with the iScript cDNA synthesis kit (Bio-Rad) and qPCR was carried out with the SsoAdvanced Universal SYBR Green Supermix (Bio-Rad). Relative expression was calculated by the 2^−(ΔΔCt)^ method using the expression of *rpoD* as the endogenous control, and the log-phase or no ATB sample as the calibrator sample. Then, the relative gene expression was plotted as the log_2_(fold change;FC) = log_2_^[2-Δ ΔCt]^.

### Statistics

Differences between the excision levels during the growth phases were assessed by one-way ANOVA followed by Dunnett’s multiple comparisons test. Differences on gene expression were assessed by two-way ANOVA followed by Dunnett’s or Tukey’s multiple comparisons test. Correlation was assessed with the Pearson correlation coefficient. All statistical analyses were carried out in GraphPad Prism v9 and v10.0.0 with α=0.05.

## ACKNOWLEDGEMENTS

This study was supported by the Millennium Institute on Immunology and Immunotherapy (ICN09_016/ICN 2021_045; former P09/016-F); grant FONDEF/ANID ID20I10082; grant FONDECYT Regular N°1231851; and grant FONDECYT Postdoctorado N°3230796.

